# SegCSR: Weakly-Supervised Cortical Surfaces Reconstruction from Brain Ribbon Segmentations

**DOI:** 10.1101/2024.12.10.626888

**Authors:** Hao Zheng, Xiaoyang Chen, Hongming Li, Tingting Chen, Peixian Liang, Yong Fan

## Abstract

Deep learning-based cortical surface reconstruction (CSR) methods heavily rely on pseudo ground truth (pGT) generated by conventional CSR pipelines as supervision, leading to dataset-specific challenges and lengthy training data preparation. We propose a new approach for reconstructing multiple cortical surfaces using *weak supervision* from brain MRI ribbon segmentations. Our approach initializes a midthickness surface and then deforms it inward and outward to form the inner (white matter) and outer (pial) cortical surfaces, respectively, by jointly learning diffeomorphic flows to align the surfaces with the boundaries of the cortical ribbon segmentation maps. Specifically, a boundary surface loss drives the initialization surface to the target inner and outer boundaries, and an inter-surface normal consistency loss regularizes the pial surface in challenging deep cortical sulci. Additional regularization terms are utilized to enforce surface smoothness and topology. Evaluated on two large-scale brain MRI datasets, our weakly-supervised method achieves comparable or superior CSR accuracy and regularity to existing supervised deep learning alternatives.

## 1 Introduction

Cortical surface reconstruction (CSR) is essential for visualizing and quantitatively analyzing cortical surfaces [1–4]. Well-established CSR pipelines like BrainSuite [5], FreeSurfer [6], and iBEAT V2.0 [7] are tailored for processing different cohorts who exhibit distinct differences in brain MRI in terms of intensity values, size, and shape, thus requiring careful parameter tuning for different datasets. Despite achieving sub-voxel accuracy and maintaining spherical topology, these pipelines involve multiple iterative steps, including surface deformation, topology checks, and corrections, leading to lengthy processing times, typically 4∼ 6h/subject.

Recently, deep learning (DL) based methods have significantly accelerated CSR by providing fast and accurate inference while preserving surface topology [8–13]. One line of research predicts implicit surface representations, such as signed distance functions [8, 14] or level sets [9], from which 3D meshes are extracted using the Marching Cubes (MC) algorithm [15] and refined with topology correction algorithms [16] to detect and rectify topology errors, ensuring that the reconstructed surface conforms to a sphere-like topology. Another line of research focuses on learning explicit surface deformations, using methods such as flow-based [11–13,17–19] or NODE-based techniques [20,21], to deform an initial mesh towards target cortical surfaces. However, DL-based CSR methods heavily depend on pseudo ground truth (pGT) surfaces generated by traditional CSR pipelines, hindering the collection of sufficiently large datasets for training and limiting generalization across diverse modalities (e.g., imaging protocols, age groups).

Given the fact that segmentation of brain structures is comparatively simpler, in this paper, we propose a ***weakly supervised*** DL framework, ***SegCSR***, to reduce the reliance on pGT in CSR via using ribbon segmentations from brain MRIs as supervision, enabling generalizing DL-based CSR approaches to scenarios where ribbon segmentation results are readily available. There are three main challenges: (1) Sub-voxel supervision signals: While existing approaches can produce precise segmentations [7, 23–26], voxel-level representations often fail to capture the fine morphology of the cerebral cortex. This is especially challenging in deep cortical sulci [6] and low-resolution images [7], where the thin, folded structure is compromised by partial volume effects (PVE) in brain MRIs. (2) Modeling surface interdependence: Reconstructing both the inner and outer cortical surfaces while maintaining their interdependent structure and spherical topology is complex [12, 19], especially without ground-truth data. This limits stability in optimizing multiple surfaces concurrently. (3) Maintaining surface topology: Ensuring mesh smoothness and topological consistency is difficult under large deformations [19, 20], as optimizing dense volumetric fields over randomly sampled vertices may distort mesh structures.

We address the diffeomorphic deformation problem in a continuous coordinate space, deforming the initialization midthickness surface towards the target inner and outer surfaces via four innovative loss functions. Specifically, the boundary surface loss function based on the ribbon segmentations and the intensity gradient loss function based on the raw image facilitate sub-voxel-level surface movement. The inter-surface normal consistency loss function explicitly integrates the normal directions of the WM, midthickness, and pial surfaces, thereby regularizing the pial surface in challenging deep cortical sulci regions. The symmetric deformation trajectory loss enforces consistency across multiple diffeomorphic deformations, explicitly coupling multiple surfaces to reduce learning complexity and regularize mesh topology.

Our main contributions are: (1) introducing a new weakly supervised paradigm for CSR, reducing dependence on pGT surfaces, (2) designing loss functions to align surfaces with cortical ribbon segmentation maps and enforce regularity of surfaces, and (3) achieving comparable or superior performance to existing DL-based CSR methods on various datasets.

## 2 Methodology

Our framework (Fig.1) reconstructs multiple cortical surfaces simultaneously and bypasses the need for conventional, time-consuming CSR-generated pGT. Section 2.1 details the network structure for coupled surface learning, while Section 2.2 illustrates loss functions that enhance sub-voxel accuracy and maintain surface topology.

**Fig. 1:**
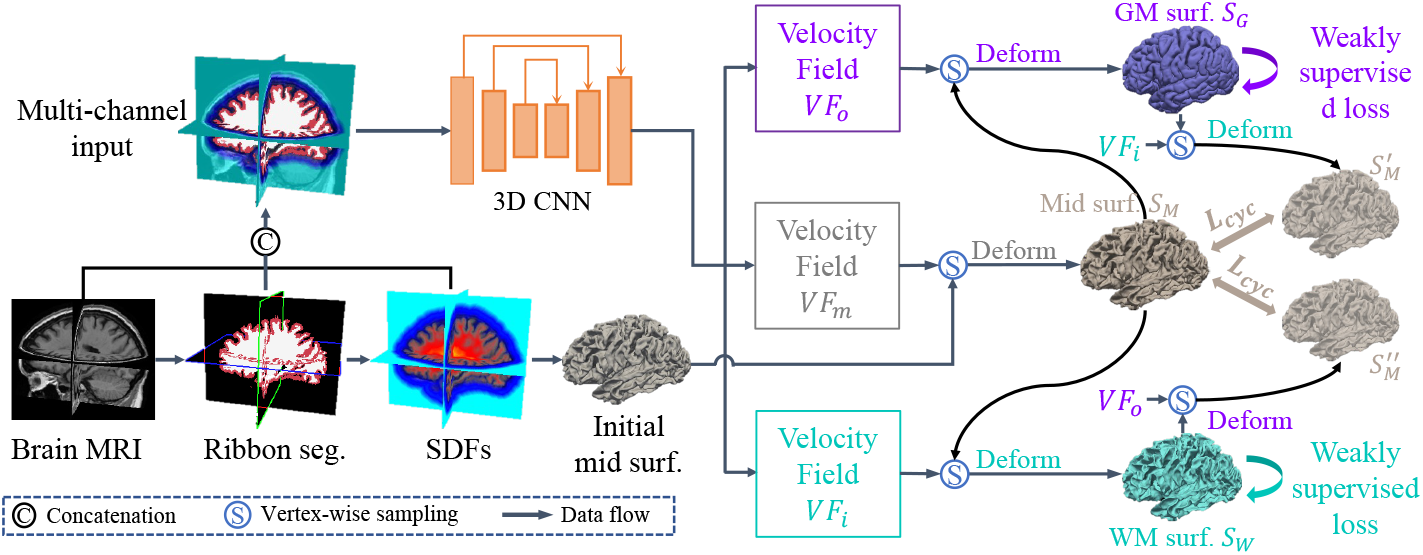
The SegCSR framework overview. SegCSR takes multi-channel images as input to learn three diffeomorphic deformations simultaneously. This optimizes the initial midthickness surface 𝒮_0_ to align with the target midthickness surface 𝒮_*M*_, then deforms 𝒮_*M*_ outwards and inwards to the pial surface 𝒮_*G*_ and WM surface 𝒮_*W*_, respectively. The model is optimized using weakly supervised loss functions: the mesh loss guides the surfaces towards the boundaries of the cortical ribbon segmentation maps; the inter-surface normal consistency loss regularizes the pial surface in deep cortical sulci; the intensity gradient loss facilitates sub-voxel-level movement; and additional regularization terms ensure deformation trajectory, topology, and smoothness.

### 2.1 Coupled Cortical Surface Reconstruction

Existing supervised methods require pGT obtained from traditional CSR pipelines to provide precise sub-voxel supervision. They can effectively learn the deformation field, even from distant initial locations, to accurately align the initialization surface with the target surfaces [11, 17, 18]. However, brain ribbon segmentation maps are inherently discrete voxel grids, offering much coarser supervision. Consequently, the selection of the initialization surface becomes more critical. Moreover, given the intricate folded patterns of the cerebral cortex, the proximity of the two banks of grooves in deep cortical sulci often poses a considerable risk of generating topology errors (e.g., handles, holes) in the reconstructed surfaces. Conversely, voxels closer to the WM surface exhibit clearer contrast, enabling a distinct separation between sulci (Fig. 2 (b)). Thus, following [19], we opt for the midthickness layer, positioned midway between the WM and pial surfaces, to serve as a connection for coupling the reconstructions of both surfaces and achieve a balanced performance for both surfaces.

**Fig. 2:**
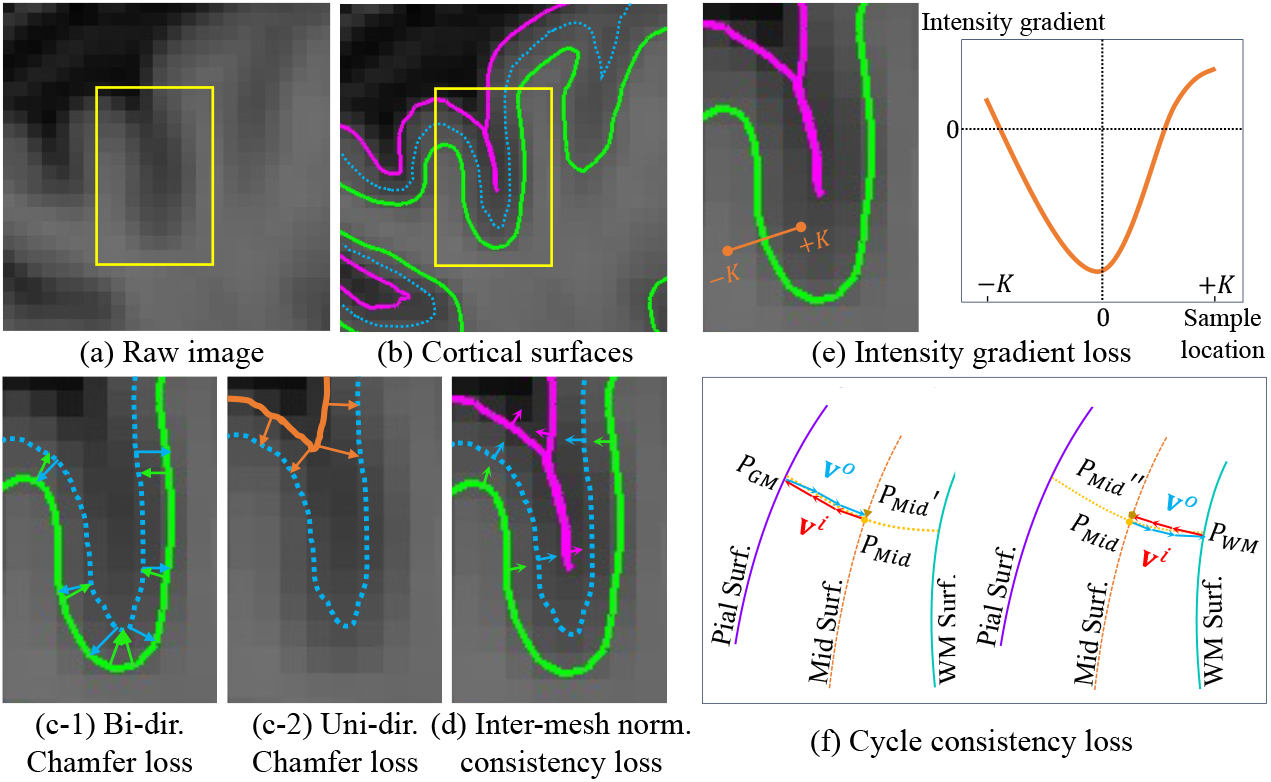
(a) Brain MRI region. (b)-(f) Loss terms illustrated: (b) WM, midthickness, pial surfaces in a deep sulcus; (c-1) Bi-directional Chamfer loss for WM surface; (c-2) Uni-directional Chamfer loss for pGT pial surface generated from the GM segmentation; (d) Normal consistency among three reconstructed surfaces; (e) Intensity gradient along a vertex normal; (f) Symmetric deformation trajectory with outward (**v**^*o*^) and inward (**v**^*i*^) velocity fields.

As illustrated in Fig. 1, SegCSR employs a neural network to jointly model three diffeomorphic flows: *F*_*θ*_(*I*, 𝒮_0_) = (**v**^*m*^, **v**^*o*^, **v**^*i*^). Here, *I* represents a multi-channel input consisting of brain MRI, cortical ribbon masks, and signed distance functions (SDFs); 𝒮_0_ denotes the initialization midthickness surface; and **v**^*m*^, **v**^*o*^, **v**^*i*^ correspond to the velocity fields that drive 𝒮_0_ towards the true midthickness surface 𝒮_*M*_, outward to the pial surface 𝒮_*G*_, and inward to the WM surface 𝒮_*W*_, respectively. The SegCSR establishes an explicit *one-to-one mapping* between multiple surfaces and is trained by minimizing weakly supervised losses between the predicted mesh and the ribbon segmentations.

The diffeomorphic deformation between the initialization surface and the target surface can be computed as the integration of an ODE [27] based on the velocity field **v**:

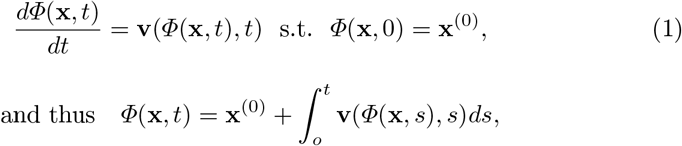

where *Φ*(**x**, *t*) defines a trajectory from the source position **x**^(0)^ = *Φ*(**x**, 0) to the target position **x**^(1)^ = *Φ*(**x**, 1). According to the *Cauchy-Lipschitz* theorem [28], if the velocity field is Lipschitz continuous, the resulting mapping *Φ*is bijective with continuous inverse. To solve this initial value problem, we perform the integration on the predicted velocity fields using standard numerical integration techniques, such as the Euler method and the Runge-Kutta method [29]. Specifically, for each integration step *t* ∈ [0, 1], each vertex’s coordinates can be updated by 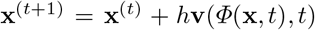, where 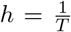 is the step size and *T* is the total time steps, and the velocity vector **v** for a vertex is trilinearly interpolated from its neighboring velocity vectors [19].

### 2.2 Weak Supervision Loss Functions

#### Mesh Loss

Weak supervision for SegCSR is derived from cortical ribbon segmentation maps of WM and GM (see Fig. 1, the filled interior area of WM and pial surfaces), which can be obtained from existing segmentation approaches [7, 23–26]. Although these ribbon segmentation maps do not perfectly represent the intricate pial surface, the WM surface is relatively easier to recognize due to its clear local intensity contrast, providing a better-separable boundary (see Fig. 2 (a-b)). Therefore, we use the boundary of the pGT WM segmentation to supervise the WM surface reconstruction. Inspired by [19, 20], we generate an SDF for the WM surface by using a distance transform algorithm, where voxels with values of zero represent the surface boundaries and voxels with negative or positive values encode their distances to the surface boundaries inward or outward, respectively. We then apply a fast topology check and correction algorithm [16] to the SDF to ensure the surface maintains spherical topology. The WM surface 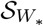 is extracted using the MC algorithm [15]. The distance of the vertices between the predicted surface 𝒮_*W*_ and the pGT surface 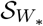 is minimized using the bi-directional Chamfer distance [11] (see Fig. 2 (c-1) for illustration):

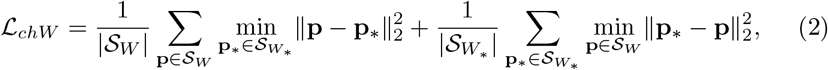

where **p** and **p**_*_ are the coordinates of vertices on meshes.

For the pial surface, GM segmentation may fail to delineate the boundary in deep cortical sulci. As shown in Fig. 2 (c-2), using a similar pGT surface generation protocol as the WM surface to generate the pial surface 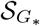 fail to capture cortical folding accurately. Directly fitting to 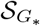 with bi-directional Chamfer loss causes the model to predict similarly inaccurate cortical sulci. To address this issue, we propose the boundary surface loss, which uses a unidirectional Chamfer distance to compute the shortest distance from the pGT pial surface 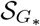 to the predicted pial surface 𝒮_*G*_:

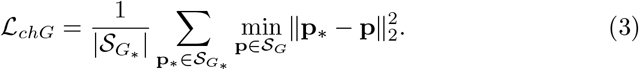

In this way, the deformed surface is not influenced by the inaccuracies of 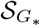 and does not move outward from the deep sulci. The overall mesh loss is computed as ℒ_*mesh*_ = ℒ_*chW*_ + ℒ_*chG*_.

#### Inter-Mesh Normal Consistency Loss

To further alleviate the difficulty of constraining the pial surface using the WM and midthickness surfaces, we propose leveraging the prior knowledge that the cerebral cortex has a sheet-like topology (i.e., the inner, middle, and outer surfaces are locally parallel to each other). As shown in Fig. 2 (d), this loss is defined to ensure that the deformation of the midthickness surface aligns with its normal direction, thereby maintaining similar normal directions on the target surfaces:

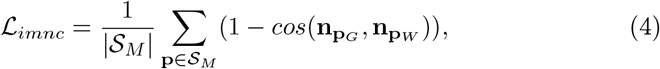

where 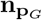 and 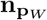 are the normal vectors of the deformed vertex **p** on 𝒮_*G*_ and 𝒮_*W*_ respectively.

#### Intensity Gradient Loss

In addition to ribbon segmentations, inspired by the fact that traditional methods utilize raw image intensity contrast to define and optimize the target surfaces, we propose to adjust the nuance between GT target surface and the pGT segmentation boundaries. By definition [6, 7], the WM (or pial) surface lies at the WM/GM (or GM/CSF) interface where image intensity change most drastically. We sample *K* points along the extended lines on each side of the normal direction at vertex **p**, and compute the gradients of neighboring points:

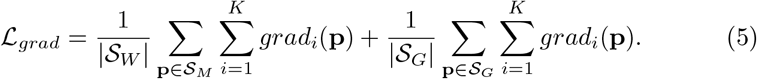

#### Cycle Consistency Loss

We utilize the midthickness layer to establish a correspondence between the inner and outer surfaces, thereby reducing the difficulty of learning large deformations. However, there is no true midthickness surface available for supervision, nor a definitive criterion for choosing between bi-directional or uni-directional approaches for different regions on the midthickness surface. Additionally, the learned velocity fields **v**^*o*^ and **v**^*i*^ could potentially cause non-inverse transformations at the midthickness surface. Inspired by [19], we devise a loss function that enforces the midthickness surface resides halfway between the WM and pial surfaces and maintains consistency along the entire trajectory:

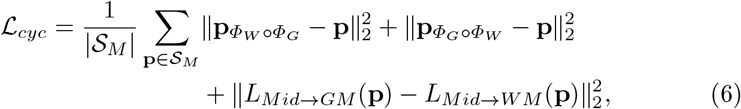

where 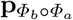 represents deforming a vertex **p** ∈ 𝒮_*M*_ with velocity field **v**^*a*^ and **v**^*b*^ sequentially, and *L*_*Mid*→*GM*_ (**p**) is the accumulated trajectory length over *T* steps of deformation. As illustrated in Fig. 2 (f), the deformations move a vertex **p**_*Mid*_ outward to **p**_*GM*_ using **v**^*o*^ and then inward to **P’**^*M id*^ using **v**^*i*^, in which the two trajectories are aligned by minimizing the distance between **p**_*M id*_ and **P’**^*M id*^. Similarly, we enforce the consistency between 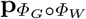 and **p**. Furthermore, starting from the midthickness layer, the trajectory lengths of the vertex moving to the WM and pial surfaces should be equal, which is regularized by the third term in Eq. 6.

In summary, we jointly optimize our SegCSR model by combining all losses: 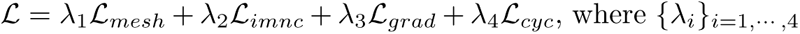 balance the loss terms.

## 3 Experiments

### 3.1 Experimental Setups

#### Datasets

We evaluate our method on two large-scale adult datasets. The ADNI-1 [30] dataset (817 subjects, ages 55-90) were randomly split into training (654), validation (50), and testing (113) subsets. The OASIS-1 [31] dataset (413 subjects, ages 18-96) were randomly split into training (330), validation (25), and testing (58) subsets The T1-weighted MRI scans were aligned to the MNI152 template and clipped to the size of 192 *×* 224 *×* 192 at 1*mm*^3^ isotropic resolution. The pseudo ground-truth (pGT) of ribbon segmentation and cortical surfaces were generated using FreeSurfer v7.2.0 [6]. The intensity values of MRI scans, ribbon segmentation maps, and SDFs were normalized to [0, 1] and the coordinates of the vertices were normalized to [− 1, 1]. Models were trained on the training sets until validation loss plateaued and tested on the test sets.

#### Implementation Details

Our framework was implemented in PyTorch and trained on a 12 GB NVIDIA P100 GPU. The 3D U-Net [22] achieved a 0.96 Dice score on ribbon segmentation. The SegCSR model used *T* = 5 steps (i.e., step size is 0.2) in an Euler solver and was optimized with Adam [32] (*β*_1_ = 0.9, *β*_2_ = 0.999, *ϵ* = 1*e*^−10^, learning rate 1*e*^−4^) for 400 epochs to reconstruct three cortical surfaces for two brain hemispheres. We set *λ*_1_ = *λ*_4_ = 1 and *λ*_2_ = *λ*_3_ = *λ*_5_ = 0.1. The surface meshes had ∼ 130*k* vertices.

#### Evaluation Metrics

To assess CSR accuracy, we used Chamfer distance (CD) [33,34], average symmetric surface distance (ASSD) [8], and 90th-percentile Hausdorff distance (HD) [8, 35], all calculated over ∼ 130*k* points uniformly sampled from the predicted and target surfaces. Topology quality was evaluated by the ratio of self-intersection faces (SIF) [8, 19, 20, 36].

### 3.2 Main Experimental Results

We compare SegCSR with representative implicit and explicit learning-based CSR approaches in Table 1 and summarize the results in Table 2.

**Table 1:**
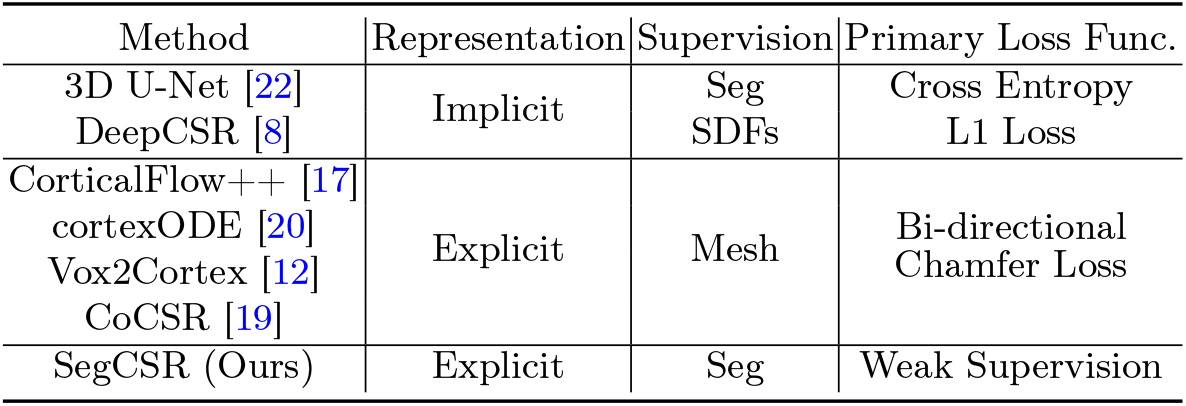
Comparison of DL-based CSR methods.

**Table 2:** Quantitative comparison with implicit and explicit learning-based methods. WM and pial surfaces were evaluated on two datasets using CD (*mm*), ASSD (*mm*), HD (*mm*), and SIF (%), reporting mean and standard deviation. Lower scores (↓) indicate better performance across metrics. “S” denotes the use of pGT surfaces from conventional pipelines, while “W” indicates weak supervision using pGT ribbon segmentations. The best results are <u>underlined</u> for the “W” supervision setting and in **bold** for “S”.

| Data | Sup. | Method | L-Pial Surface |  |  |  |
| --- | --- | --- | --- | --- | --- | --- |
| | | | CD ↓ | ASSD ↓ | HD ↓ | SIF ↓ |
| ADNI | S | Vox2Cortex [12] | 0.582±0.028 | 0.370±0.025 | 0.746±0.057 | 0.059±0.039 |
| | | CorticalFlow++ [17] | 0.545±0.036 | 0.410±0.033 | 0.886±0.069 | 0.098±0.067 |
| | | cortexODE [20] | 0.476±0.017 | 0.214±0.020 | 0.455±0.058 | <u>0.022±0.012</u> |
| | W | DeepCSR [8] | 0.945±0.078 | 0.593±0.065 | 1.149±0.203 | \ |
| | | 3D U-Net [22] | 0.598±0.049 | 0.341±0.037 | 0.782±0.163 | \ |
| | | SegCSR (Ours) | **0.576±0.019** | **0.323±0.019** | **0.747±0.046** | 0.012±0.011 |
| OASIS | S | Vox2Cortex [12] | 0.588±0.032 | 0.381±0.030 | 0.750±0.063 | 0.061±0.037 |
| | | CorticalFlow++ [17] | 0.531±0.035 | 0.399±0.030 | 0.812±0.057 | 0.088±0.045 |
|  |  | cortexODE [20] | <u>0.481±0.019</u> | <u>0.218±0.021</u> | <u>0.461±0.062</u> | <u>0.026±0.015</u> |
| | W | DeepCSR [8] | 0.986±0.085 | 0.617±0.070 | 1.331±0.212 | \ |
| | | 3D U-Net [22] | 0.611±0.069 | 0.332±0.050 | 0.774±0.267 | \ |
| | | SegCSR (Ours) | **0.581±0.016** | **0.320±0.019** | **0.723±0.038** | 0.013±0.012 |
| Data | Sup. | Method | L-WM Surface |  |  |  |
| | | | CD ↓ | ASSD ↓ | HD ↓ | SIF ↓ |
| ADNI | S | Vox2Cortex [12] | 0.577±0.027 | 0.353±0.022 | 0.722±0.055 | 0.043±0.023 |
| | | CorticalFlow++ [17] | 0.544±0.034 | 0.401±0.030 | 0.878±0.066 | 0.069±0.042 |
|  |  | cortexODE [20] | <u>0.458±0.016</u> | <u>0.192±0.015</u> | <u>0.436±0.014</u> | <u>0.015±0.011</u> |
| | W | DeepCSR [8] | 0.938±0.076 | 0.587±0.064 | 1.137±0.193 | \ |
| | | 3D U-Net [22] | 0.473±0.013 | 0.265±0.015 | 0.558±0.028 | \ |
| | | SegCSR (Ours) | **0.467±0.015** | **0.257±0.020** | **0.542±0.036** | 0.011±0.011 |
| OASIS | S | Vox2Cortex [12] | 0.581±0.028 | 0.375±0.027 | 0.731±0.059 | 0.046±0.027 |
| | | CorticalFlow++ [17] | 0.529±0.033 | 0.398±0.030 | 0.810±0.055 | 0.086±0.042 |
|  |  | cortexODE [20] | <u>0.463±0.018</u> | <u>0.207±0.017</u> | <u>0.435±0.015</u> | <u>0.018±0.010</u> |
| | W | DeepCSR [8] | 0.975±0.081 | 0.594±0.067 | 1.151±0.197 | \ |
| | | 3D U-Net [22] | 0.454±0.013 | 0.245±0.017 | 0.489±0.031 | \ |
| | | SegCSR (Ours) | **0.449±0.011** | **0.222±0.016** | **0.460±0.026** | 0.013±0.011 |

I. ***Comparison with Implicit Approaches***. 3D U-Net [22] generates ribbon segmentation maps and DeepCSR [8] outputs SDFs encoding cortical surfaces. Post-processing is required to correct topology and extract a mesh, resulting in SIFs of 0. Otherwise, the SIFs for 3D U-Net’s WM and pial surfaces range from 3% to 15%. SegCSR achieves superior geometric accuracy and produces a negligible number of self-intersecting faces, ∼ 0.3% on average for both white and pial surfaces. Fig. 3 shows that SegCSR effectively deforms the pial surface into deep sulci, while the baseline approaches exhibit large geometric errors due to the PVE problem of brain MRI. Additionally, SegCSR requires only 0.37s of runtime per brain hemisphere, orders of magnitude faster than the traditional FreeSurfer pipeline.

**Fig. 3:**
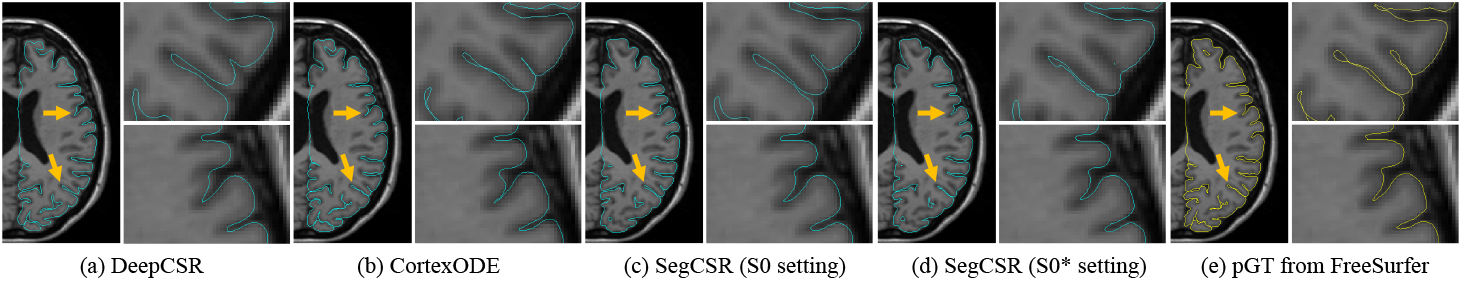
Qualitative comparison with DeepCSR and CortexODE.
II. ***Comparison with Explicit Approaches***. Explicit learning-based methods [12, 17, 20] utilize pGT surfaces from conventional pipelines, providing higher-precision supervision compared to the pGT ribbon segmentations used in our SegCSR. Although SegCSR performs below the top performer cortexODE [20] (e.g., 17% and 2% lower in CD for the left pial and WM surfaces, respectively), it surpasses other supervised methods in both geometric and morphological accuracy, outperforming [12] by 1% and 19% in CD for the left pial and WM surfaces. These results demonstrate SegCSR’s strong potential as a viable alternative when precise surface supervision is unavailable.

### 3.3 Ablation Studies

We evaluated the contribution of different losses of our method to the surface reconstruction performance in terms of both accuracy (CD, ASSD, HD) and topological correctness (SIF). The results are summarized in Table 3. **(I)** The setting S3 represents using our proposed Chamfer loss (i.e., uni-directional for the pial surface) alone, while S3^⋆^ referes to using existing bi-directional Chamfer loss for both WM and pial surfaces. The results of S3 and S3^⋆^ indicated that the model using bi-directional Chamfer loss overfitted to the pGT segmentation boundary and failed to fit the deep cortical sulci. Another pair of comparison, S0 and S0^⋆^, showed a similar phenomenon. **(II)** Enforcing the inter-mesh normal consistency of the WM and pial surfaces (S2, ℒ_*imnc*_) improved geometric accuracy by explicitly constraining the normal direction of two surfaces but slightly worsened the topology, which might be caused by the discrepancy between the midthickness and the WM (and pial) surface. **(III)** The proposed intensity gradient loss (S1, ℒ_*grad*_) helped adjust the deformed surfaces locally, leading to slightly improved geometric accuracy and reduced topology error. **(IV)** Enforcing equality of the trajectories from the midthickness surface to the WM and pial surfaces and symmetric cycle consistency of two trajectories (S0, ℒ_*cyc*_) helped optimize the midthickness surface and promoted the invertibility of deformations. Overall, our proposed method struck a balance between geometric accuracy and topology quality, with each component playing a complementary role.

**Table 3:** Ablation studies on the impact of loss terms on the ADNI dataset. The setting S0 refers to our complete setting (cf. Table 2).

|  | Loss |  |  |  | L-Pial Surface |  |  |  | L-WM Surface |  |  |  |
| --- | --- | --- | --- | --- | --- | --- | --- | --- | --- | --- | --- | --- |
| | $\mathcal{L}_{mesh}$ | $\mathcal{L}_{imnc}$ | $\mathcal{L}_{grad}$ | $\mathcal{L}_{cyc}$ | CD | ASSD | HD | SIF | CD | ASSD | HD | SIF |
| S0 | ✓ | ✓ | ✓ | ✓ | 0.576 | 0.323 | 0.747 | 0.012 | 0.467 | 0.257 | 0.542 | 0.011 |
| S1 | ✓ | ✓ | ✓ |  | 0.579 | 0.325 | 0.748 | 0.014 | 0.469 | 0.248 | 0.544 | 0.015 |
| S2 | ✓ | ✓ |  |  | 0.579 | 0.325 | 0.749 | 0.018 | 0.473 | 0.249 | 0.544 | 0.017 |
| S3 | ✓ |  |  |  | 0.589 | 0.356 | 0.764 | 0.015 | 0.473 | 0.256 | 0.564 | 0.014 |
| S0* | ✓* | ✓ | ✓ | ✓ | 0.606 | 0.325 | 0.749 | 0.030 | 0.469 | 0.257 | 0.545 | 0.026 |
| S3* | ✓* |  |  |  | 0.626 | 0.321 | 0.773 | 0.034 | 0.476 | 0.256 | 0.562 | 0.031 |

## 4 Conclusions

We introduce SegCSR, a novel approach to jointly reconstruct multiple cortical surfaces using weak supervision from ribbon segmentations derived from brain MRIs. Our method initializes a midthickness surface and then deforms it inward and outward to the inner and outer cortical surfaces by jointly learning diffeomorphic flows. The new boundary loss function optimizes the surfaces toward the boundaries of the cortical ribbon segmentation maps while the inter-surface normal consistency loss regularizes the pial surface in complex and challenging cortical sulci regions. Additional regularization terms are incorporated to enforce reconstructed surfaces’ smoothness and topology. Extensive experiments conducted on two large-scale brain MRI datasets demonstrate superior performance in terms of accuracy and surface regularity compared to existing supervised DL-based alternatives.

## 5 Compliance with ethical standards

This research was conducted retrospectively using human subject data made available in open access [30, 31]. Ethical approval was not required as confirmed by the license attached with the public data.

## 6 Acknowledgments

This work was supported in part by the NIH grants AG066650 and EB022573.

## References

1. D. C. Van Essen, H. A. Drury, S. Joshi, and M. I. Miller, “Functional and structural mapping of human cerebral cortex: solutions are in the surfaces,” PNAS, vol. 95, no. 3, pp. 788–795, 1998.

2. A. M. Dale, B. Fischl, and M. I. Sereno, “Cortical surface-based analysis: I. segmentation and surface reconstruction,” Neuroimage, vol. 9, no. 2, pp. 179–194, 1999.

3. J. M. Roe, D. Vidal-Piñeiro, Ø. Sørensen, A. M. Brandmaier, S. Düzel, H. A. Gonzalez, R. A. Kievit, E. Knights, S. Kühn, U. Lindenberger, et al., “Asymmetric thinning of the cerebral cortex across the adult lifespan is accelerated in alzheimer’s disease,” Nat. Commun., vol. 12, no. 1, p. 721, 2021.

4. L. M. Rimol, R. Nesvåg, D. J. Hagler Jr, Ø. Bergmann, C. Fennema-Notestine, C. B. Hartberg, U. K. Haukvik, E. Lange, C. J. Pung, A. Server, et al., “Cortical volume, surface area, and thickness in schizophrenia and bipolar disorder,” Biological Psychiatry, vol. 71, no. 6, pp. 552–560, 2012.

5. D. W. Shattuck and R. M. Leahy, “Brainsuite: an automated cortical surface identification tool,” MedIA, vol. 6, no. 2, pp. 129–142, 2002.

6. B. Fischl, “Freesurfer,” Neuroimage, vol. 62, no. 2, pp. 774–781, 2012.

7. L. Wang, Z. Wu, L. Chen, Y. Sun, W. Lin, and G. Li, “ibeat v2.0: a multisite-applicable, deep learning-based pipeline for infant cerebral cortical surface reconstruction,” Nat Protocols, vol. 18, no. 5, pp. 1488–1509, 2023.

8. R. S. Cruz, L. Lebrat, P. Bourgeat, C. Fookes, J. Fripp, and O. Salvado, “DeepCSR: A 3D deep learning approach for cortical surface reconstruction,” in WACV, pp. 806–815, 2021.

9. J. Ren, Q. Hu, W. Wang, W. Zhang, C. S. Hubbard, P. Zhang, N. An, Y. Zhou, L. Dahmani, D. Wang, et al., “Fast cortical surface reconstruction from MRI using deep learning,” Brain Inform., vol. 9, no. 1, pp. 1–16, 2022.

10. Q. Ma, E. C. Robinson, B. Kainz, D. Rueckert, and A. Alansary, “PialNN: A fast deep learning framework for cortical pial surface reconstruction,” in IWMLCN, pp. 73–81, Springer, 2021.

11. L. Lebrat, R. Santa Cruz, F. de Gournay, D. Fu, P. Bourgeat, J. Fripp, C. Fookes, and O. Salvado, “CorticalFlow: A diffeomorphic mesh transformer network for cortical surface reconstruction,” NeurIPS, vol. 34, pp. 29491–29505, 2021.

12. F. Bongratz, A.-M. Rickmann, S. Pölsterl, and C. Wachinger, “Vox2Cortex: Fast explicit reconstruction of cortical surfaces from 3D MRI scans with geometric deep neural networks,” in CVPR, pp. 20773–20783, 2022.

13. A. Hoopes, J. E. Iglesias, B. Fischl, D. Greve, and A. V. Dalca, “Topofit: Rapid reconstruction of topologically-correct cortical surfaces,” in MIDL, 2022.

14. Y. Hong, S. Ahmad, Y. Wu, S. Liu, and P.-T. Yap, “Vox2Surf: Implicit surface reconstruction from volumetric data,” in IWMLMI, pp. 644–653, Springer, 2021.

15. T. Lewiner, H. Lopes, A. W. Vieira, and G. Tavares, “Efficient implementation of marching cubes’ cases with topological guarantees,” Journal of Graphics Tools, vol. 8, no. 2, pp. 1–15, 2003.

16. P.-L. Bazin and D. L. Pham, “Topology correction of segmented medical images using a fast marching algorithm,” CMPB, vol. 88, no. 2, pp. 182–190, 2007.

17. R. Santa Cruz, L. Lebrat, D. Fu, P. Bourgeat, J. Fripp, C. Fookes, and O. Salvado, “CorticalFlow++: Boosting cortical surface reconstruction accuracy, regularity, and interoperability,” in MICCAI, pp. 496–505, Springer, 2022.

18. X. Chen, J. Zhao, S. Liu, S. Ahmad, and P.-T. Yap, “SurfFlow: A flow-based approach for rapid and accurate cortical surface reconstruction from infant brain mri,” in MICCAI, pp. 380–388, Springer, 2023.

19. H. Zheng, H. Li, and Y. Fan, “Coupled reconstruction of cortical surfaces by diffeomorphic mesh deformation,” NeurIPS, vol. 37, 2023.

20. Q. Ma, L. Li, E. C. Robinson, B. Kainz, D. Rueckert, and A. Alansary, “CortexODE: Learning cortical surface reconstruction by neural ODEs,” TMI, vol. 42, no. 2, pp. 430–443, 2022.

21. Q. Ma, L. Li, V. Kyriakopoulou, J. V. Hajnal, E. C. Robinson, B. Kainz, and D. Rueckert, “Conditional temporal attention networks for neonatal cortical surface reconstruction,” in MICCAI, pp. 312–322, Springer, 2023.

22. O. Ronneberger, P. Fischer, and T. Brox, “U-Net: Convolutional networks for biomedical image segmentation,” in MICCAI, pp. 234–241, 2015.

23. A. G. Roy, S. Conjeti, N. Navab, C. Wachinger, A. D. N. Initiative, et al., “Quicknat: A fully convolutional network for quick and accurate segmentation of neuroanatomy,” NeuroImage, vol. 186, pp. 713–727, 2019.

24. L. Henschel, S. Conjeti, S. Estrada, K. Diers, B. Fischl, and M. Reuter, “Fastsurfer-a fast and accurate deep learning based neuroimaging pipeline,” NeuroImage, vol. 219, p. 117012, 2020.

25. Y. Li, H. Li, and Y. Fan, “ACEnet: Anatomical context-encoding network for neuroanatomy segmentation,” MedIA, vol. 70, p. 101991, 2021.

26. B. Billot, D. N. Greve, O. Puonti, A. Thielscher, K. Van Leemput, B. Fischl, A. V. Dalca, J. E. Iglesias, et al., “Synthseg: Segmentation of brain mri scans of any contrast and resolution without retraining,” MedIA, vol. 86, p. 102789, 2023.

27. V. I. Arnold, Ordinary differential equations. Springer Science & Business Media, 1992.

28. G. Teschl, Ordinary differential equations and dynamical systems, vol. 140. American Mathematical Soc., 2012.

29. R. L. Burden, J. D. Faires, and A. M. Burden, Numerical analysis. Cengage learning, 2015.

30. C. R. Jack Jr, M. A. Bernstein, N. C. Fox, P. Thompson, G. Alexander, D. Harvey, B. Borowski, P. J. Britson, J. L. Whitwell, C. Ward, et al., “The alzheimer’s disease neuroimaging initiative (ADNI): MRI methods,” J. Magn. Reson. Imaging, vol. 27, no. 4, pp. 685–691, 2008.

31. D. S. Marcus, T. H. Wang, J. Parker, J. G. Csernansky, J. C. Morris, and R. L. Buckner, “Open access series of imaging studies (OASIS): cross-sectional MRI data in young, middle aged, nondemented, and demented older adults,” J. Cogn. Neurosci., vol. 19, no. 9, pp. 1498–1507, 2007.

32. D. P. Kingma and J. Ba, “Adam: A method for stochastic optimization,” in ICLR, 2015.

33. H. Fan, H. Su, and L. J. Guibas, “A point set generation network for 3d object reconstruction from a single image,” in CVPR, pp. 605–613, 2017.

34. N. Wang, Y. Zhang, Z. Li, Y. Fu, H. Yu, W. Liu, X. Xue, and Y.-G. Jiang, “Pixel2Mesh: 3D mesh model generation via image guided deformation,” TPAMI, vol. 43, no. 10, pp. 3600–3613, 2020.

35. A. A. Taha and A. Hanbury, “Metrics for evaluating 3D medical image segmentation: analysis, selection, and tool,” BMC Med. Imaging, vol. 15, no. 1, pp. 1–28, 2015.

36. R. Dahnke, R. A. Yotter, and C. Gaser, “Cortical thickness and central surface estimation,” Neuroimage, vol. 65, pp. 336–348, 2013.

